# Interactions between RNA m^6^A modification, alternative splicing, and poly(A) tail revealed by MePAIso-seq2

**DOI:** 10.1101/2021.08.29.458071

**Authors:** Yusheng Liu, Yiwei Zhang, Falong Lu, Jiaqiang Wang

## Abstract

RNA post-transcriptional regulation involves 5′-end capping, 3′ poly(A) tailing (including polyadenylation sites, tail length, and non-A residues^1, 2^), alternative splicing, and chemical modifications including N^6^-methyladenosine (m^6^A)^3^. Studying the interplay of m^6^A, alternative splicing, alternative polyadenylation sites, poly(A) tail length, and non-A residues in poly(A) tails requires monitoring them simultaneously on one transcript, however strategies to achieve this are lacking. Therefore, we developed a new method, combining m^6^A-specific methylated RNA immunoprecipitation and the PacBio-based, tail-included, full-length RNA sequencing approach PAIso-seq2, which we’ve named m^6^A and poly(A) inclusive RNA isoform sequencing 2 (MePAIso-seq2). Using MePAIso-seq2, we revealed that m^6^A promotes and inhibits a similar number of alternative splicing events in mouse cell lines, showing that m^6^A does affect alternative splicing. In contrast, no correlation was detected between m^6^A and alternative polyadenylation sites choice. Surprisingly, we found that m^6^A-modified RNAs possess longer poly(A) tails and a lower proportion of poly(A) tails containing non-A residues, especially in mouse embryonic stem cells. Together, we developed a new method to detect full-length m^6^A-modified RNAs to comprehensively study the relationships between m^6^A, alternative splicing, and poly(A) tailing, laying a foundation for further exploration of the functional coordination of different RNA post-transcriptional modifications.

## Introduction

N^6^-methyladenosine (m^6^A) has been recognized as a post-transcriptional RNA chemical modification in mammalian cells since 1974^4^. With the rapid development of immunoprecipitation and high throughput sequencing, m^6^A has been found to be the most common type of chemical modification in mammalian RNA, including mRNAs and long non-coding RNAs (lncRNAs)^5–7^. m^6^A has functions in a wide array of physiological and biological processes, such as self-renewal and differentiation of embryonic stem cells^8^, T cell homeostasis^9^, ischemic heart failure^10, 11^, adipogenesis^12^, cancer^13, 14^, spermatogenesis^15^, and circadian clock rhythms^16^. In both mammals and yeast, RNA m^6^A preferentially occurs in both gene coding regions and 3′ untranslated regions (3′-UTRs), implicating its fundamental roles in every aspect of post-transcriptional regulation, including splicing^12^, subcellular transport^17^, decay^18^, and translation^19, 20^.

Alternative splicing is a highly dynamic and complex process that greatly increases RNA and protein biodiversity by allowing a single gene to encode for multiple proteins or different structural non-coding RNAs, influencing most biological functions in eukaryotes. Alternative splicing occurs as a conserved phenomenon in eukaryotes, and in humans about 95% of multi-exonic genes are alternatively spliced^21^. Recently alternative splicing was also identified in adenovirus type 2, and was found to increase the variability of the virus^22^. The poly(A) tails are homopolymeric sequences at the 3′-ends of RNA molecules, and has long been recognized as an essential post-transcriptional modification of most mRNAs and lncRNAs. The presence of a poly(A) tail is required for mRNA nuclear transport, stability, and translation^2, 23–25^. The regulation through poly(A) tail length can control localized mRNA translation in diverse biological processes, including immune response^26^, learning and memory^27–29^, and germ cell development^25, 30^. In addition to the length, non-A residues in poly(A) tails (non-A residues for short hereafter) are wide-spread in both the 3′-ends and internal positions^1, 2, 31^. The 3′-end U residues promote mRNA degradation, while the 3′-end G residues stabilizes the mRNA^32, 33^. The biological importance of these non-A residues mediated regulation has been demonstrated in the oocyte-to-embryo transition (OET) in mice, rats, pigs and humans^34–38^.

m^6^A modifications, alternative splicing, and poly(A) tailing constitute multiple layers of post-transcriptional RNA regulation, expanding the RNA complexity from a single gene. Currently, if and how cross-talk occurs between these different post-transcriptional modification processes is unknown and requires in depth study. However, there are currently no strategies available to achieve the analysis of all these modifications simultaneously on a single transcript. Current m^6^A mapping methods based on short-reads sequencing are incapable of capturing the full-length transcript and poly(A) tail sequence information. PAIso-seq1^2^, PAIso-seq2^39^, and FLAM-seq^1^ can provide a poly(A) tail inclusive full-length transcript sequence, but cannot detect m^6^A modifications, while Nanopore native RNA sequencing can not detect non-A residues in poly(A) tails due to inability in sequencing homopolymers^40, 41^.

Here, we combined m^6^A RNA immunoprecipitation (RIP) and PAIso-seq2, which provides poly(A) tail inclusive full length cDNA sequences, to achieve a new method called m^6^A and poly(A) inclusive RNA isoform sequencing 2 (MePAIso-seq2), that can be used to investigate interplay between m^6^A, alternative splicing, poly(A) tail length, and non-A residues.

## Results

### Establishment of MePAIso-seq2

By combining m^6^A RIP and PAIso-seq2 methods, we developed a new method, named MePAIso-seq2, for analyzing m^6^A-modified poly(A) inclusive full-length transcripts (Fig. 1a). This method employs adaptor ligation to preserve RNA tails regardless of tails polyadenylated or non-polyadenylated. We performed MePAIso-seq2 using two different previously validated antibodies for m^6^A RIP (m^6^A-1 and m^6^A-2) in NIH 3T3 cells (3T3) and in mouse embryonic stem cells (ES). The results showed that the reproducibility of MePAIso-seq2 was very good, not only between two duplicates but also between the two antibodies (Fig. 1b and Extended Data Fig. 1). We compared our data to that from the traditional m^6^A mapping method MeRIP-Seq^42, 43^. High correlations between our data and others were observed (Fig. 1c). These results indicate that MePAIso-seq2 can sequence and analyze m^6^A-modified full-length transcripts accurately and reproducibly.

**Fig. 1.**
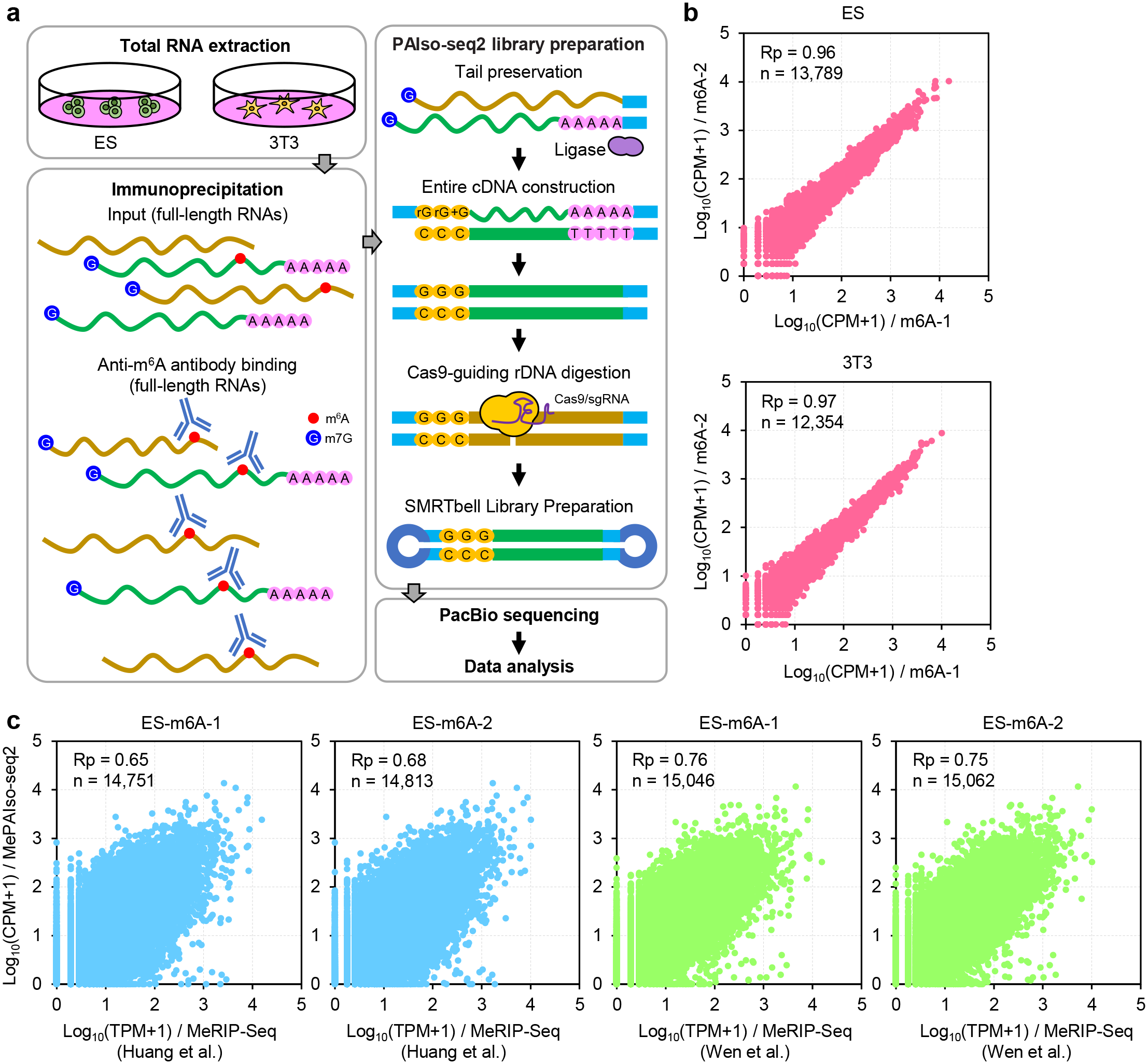
MePAIso-seq2 for full-length poly(A) inclusive sequencing of m^6^A modified RNA. **a,** Flowchart for MePAIso-seq2, which contains m^6^A modified RNA immunoprecipitation (RIP), PAIso-seq2 library construction, and PacBio sequencing. **b,** Scatter plot showing the Pearson correlation of gene expression between two m^6^A RIP antibodies (m^6^A-1, Active Motif, #91261; m^6^A-2, Abcam, #ab151230) in log10 scale in 3T3 and ES cell lines. Each dot represents one gene. Pearson’s correlation coefficient (Rp) and number of genes included in the analysis are shown at the top left of each graph. **c,** Scatter plot showing the Pearson correlation of gene expression between MePAIso-seq2 and MeRIP-Seq^42, 43^ in log10 scale in 3T3 and ES cell lines. Each dot represents one gene. Pearson’s correlation coefficient (Rp) and number of genes included in the analysis are shown at the top left of each graph.

### m^6^A regulates mRNA splicing

Some observations suggest a role of m^6^A in RNA splicing—especially skipped exon (SE) inhibition^12, 15, 44^—by RNA-seq analysis in samples depletion of METTL3^5, 15^ (methyltransferase like 3, m^6^A writer^45^), FTO^12^ (fat-mass and obesity-associated protein, m^6^A eraser), or ALKBH5^44^ (α-ketoglutarate-dependent dioxygenase alkB homolog 5, m^6^A eraser). However, it is also reported that m^6^A may not be required for splicing but does regulate cytoplasmic mRNA turnover^46, 47^. One advantage of MePAIso-seq2 is its ability to sequence full-length transcripts, therefore allowing accurate study of the relationship between m^6^A and alternative splicing (AS).

We calculated the number of seven types of AS events, including alternative 3′ splice site (A3), alternative 5′ splice site (A5), alternative first exon (AF), alternative last exon (AL), mutually exclusive exon (MX), retained intron (RI), and SE (Fig. 2a). We used SUPPA2 to compare m^6^A RIP samples to input samples and calculated the ΔPSI (percent spliced in index (PSI) of input minus PSI of m^6^A RIP for each AS event, described in the ‘Methods’ section). When there are multiple m^6^A sites and multiple AS event in one transcript, the calculated ΔPSI must be less than the theoretical value because each m^6^A may not affect all AS events (Extended Data Fig. 2a). So, when the ΔPSI value for one AS event is not zero and statistically significant, the AS event is thought to be regulated by m^6^A. We identified 597 and 1,017 AS events, which were regulated by m^6^A (ΔPSI ≠ 0, *p < 0.05*) in 3T3 and ES, respectively (Fig. 2b, left).

**Fig. 2.**
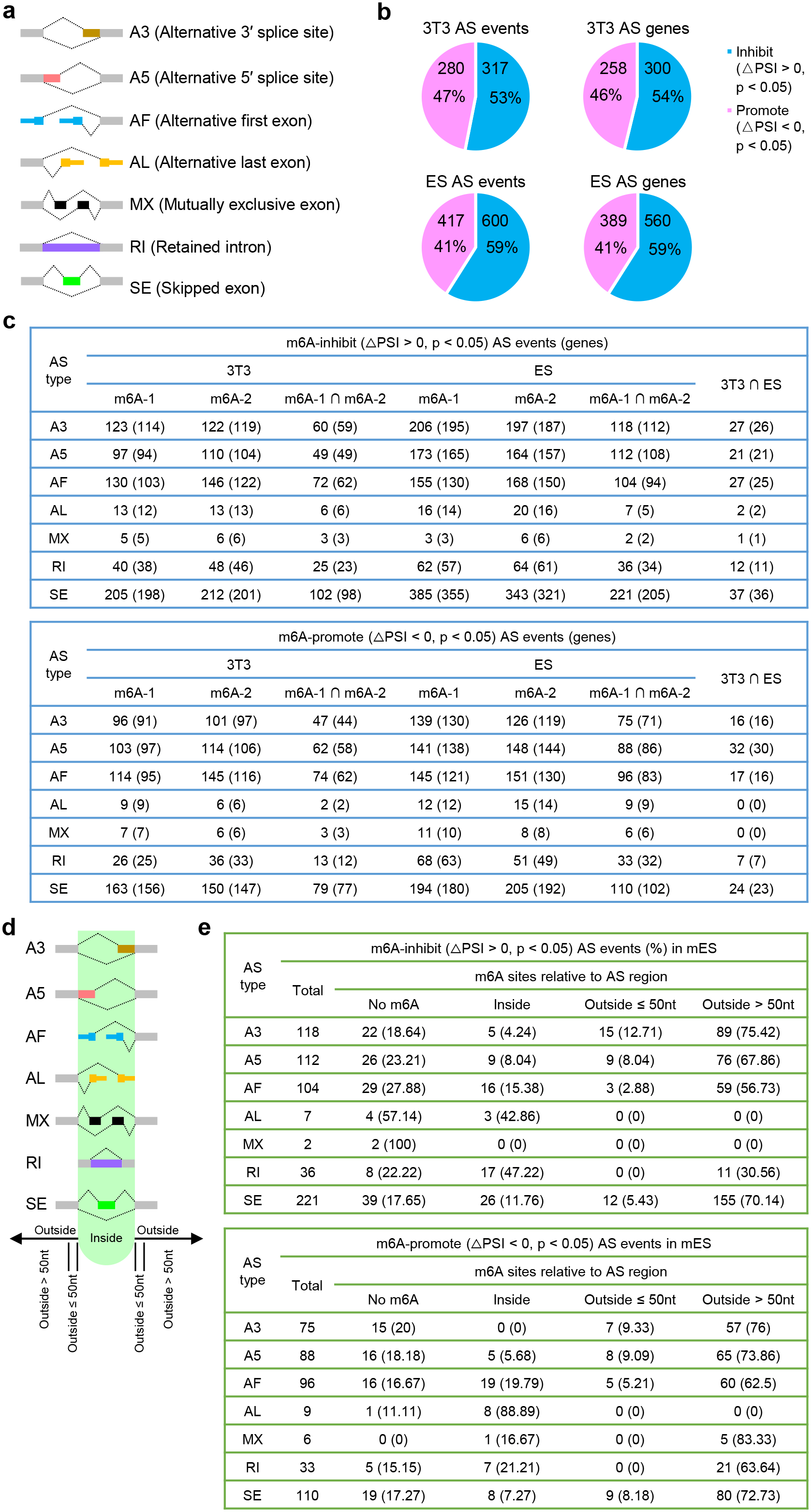
m^6^A regulates mRNA alternative splicing. **a,** Schematic diagram of the seven types of AS events. AS, alternative splicing; A5, alternative 5′ splice site; A3, alternative 3′ splice site; AF, alternative first exon; AL, alternative last exon; MX, mutually exclusive exon; RI, retained intron; SE, skipped exon. **b,** Counts of AS events and covered genes inhibited or promoted by m^6^A in 3T3 and ES cell lines. **c,** Tables for the counts of m^6^A-inhibition (up) and m^6^A-promotion (down) AS events and AS genes of the seven AS types in 3T3 and ES cell lines. **d,** Model for classification of m^6^A sites into three groups based on location relative to the AS region (green). **e,** Tables for the counts of m^6^A-inhibition (up) and m^6^A-promotion (down) of the seven types of AS events relative to AS region (**d**) in ES. Locations of m^6^A sites relative to the alternative splicing region are calculated from the data of others^48^. No m^6^A means there is no m^6^A site in the transcript according to the reference^48^. Using SUPPA2 to calculate the percent spliced in index (PSI) of each AS event with the default parameters. ΔPSI, PSI of input minus PSI of m^6^A RIP samples. ∩, intersection.

We also examined the number of AS genes associated with each type of AS event and found that the AS of 558 and 949 genes were regulated by m^6^A in 3T3 and ES, respectively (Fig. 2b, right). Additionally, we can see that the proportion of m^6^A-promotion (ΔPSI < 0, *p < 0.05*) and m^6^A-inhibition (ΔPSI > 0, *p < 0.05*) is similar in 3T3, for both AS events and AS genes (Fig. 2b, up). While more m^6^A-inhibition (59%) than m^6^A-promotion (41%) was found in ES, for both AS events and AS genes (Fig. 2b, down). Listed in detail are the counts of m^6^A-promotion (ΔPSI < 0, *p < 0.05*) and m^6^A-inhibition (ΔPSI > 0, *p* < 0.05) AS events and AS genes corresponding to the seven types of AS (Fig. 2c). The proportions of the seven types of AS events identified from m^6^A RIP data distributed similarly to that of the input (Fig. 2c and Extended Data Fig. 2b), indicating that m^6^A regulates AS with no obvious bias in regard to AS type.

Since it is reported that m^6^As are significantly enriched in alternatively spliced exons^5^, we tested this finding using our long-read data. The precise m^6^A sites within the transcripts in 3T3 and ES were obtained from the m^6^A-atlas^48^. We classified m^6^A sites into three groups based on location relative to the AS region (Fig. 2d), and listed the counts of m^6^A-promotion (ΔPSI < 0, *p < 0.05*) and m^6^A-inhibition (ΔPSI > 0, *p* < 0.05) AS events for the seven AS types in ES (Fig. 2e). The results showed that m^6^A sites predominantly located more than 50 nt outside of the AS region (Fig. 2e). We conclude that m^6^A does affect AS events, where both promotion and inhibition are involved for the different AS event types.

### Relationship between m^6^A and alternative polyadenylation sites

The ability to sequence transcripts including full-length poly(A) tails using MePAIso-seq2 allows us to study the relationship between m^6^A and poly(A) tailing. It is reported that m^6^A is predominantly (≥ 70%) located in the last exon allowing for potential regulation of PAS choice^49^, therefore we investigated the effects of m^6^A on PAS choice (Fig. 3a). The results showed that m^6^A modified mRNAs slightly prefers the transcripts with proximal PAS (pPAS) over transcripts with distal PAS (dPAS) in 3T3 cells (Fig. 3b), while minimal preference was seen in ES cells (Fig. 3d). These results indicate that m^6^A modifications have very slight preference on PAS choice and is regulated differently in 3T3 and ES cells. It is reported that there is a statistically significant trend for isoforms using dPAS to carry longer poly(A) tails than isoforms using pPAS in Hela cells and human induced pluripotent stem cells^1^. Thus, we wondered whether m^6^A affects poly(A) tail length for different isoforms using pPAS or dPAS. We computed the tail length distributions for isoforms using pPAS or dPAS and found no obvious consistent difference on the distribution of poly(A) tail length between isoforms using pPAS and isoforms using dPAS for both input and m^6^A RIP samples in 3T3 cells (Fig. 3c). In ES cells, the poly(A) tails on isoforms using dPAS were significantly longer (Fig. 3e). These results indicate that the effect of m^6^A on poly(A) tailing is differentially regulated in 3T3 and ES cells.

**Fig. 3.**
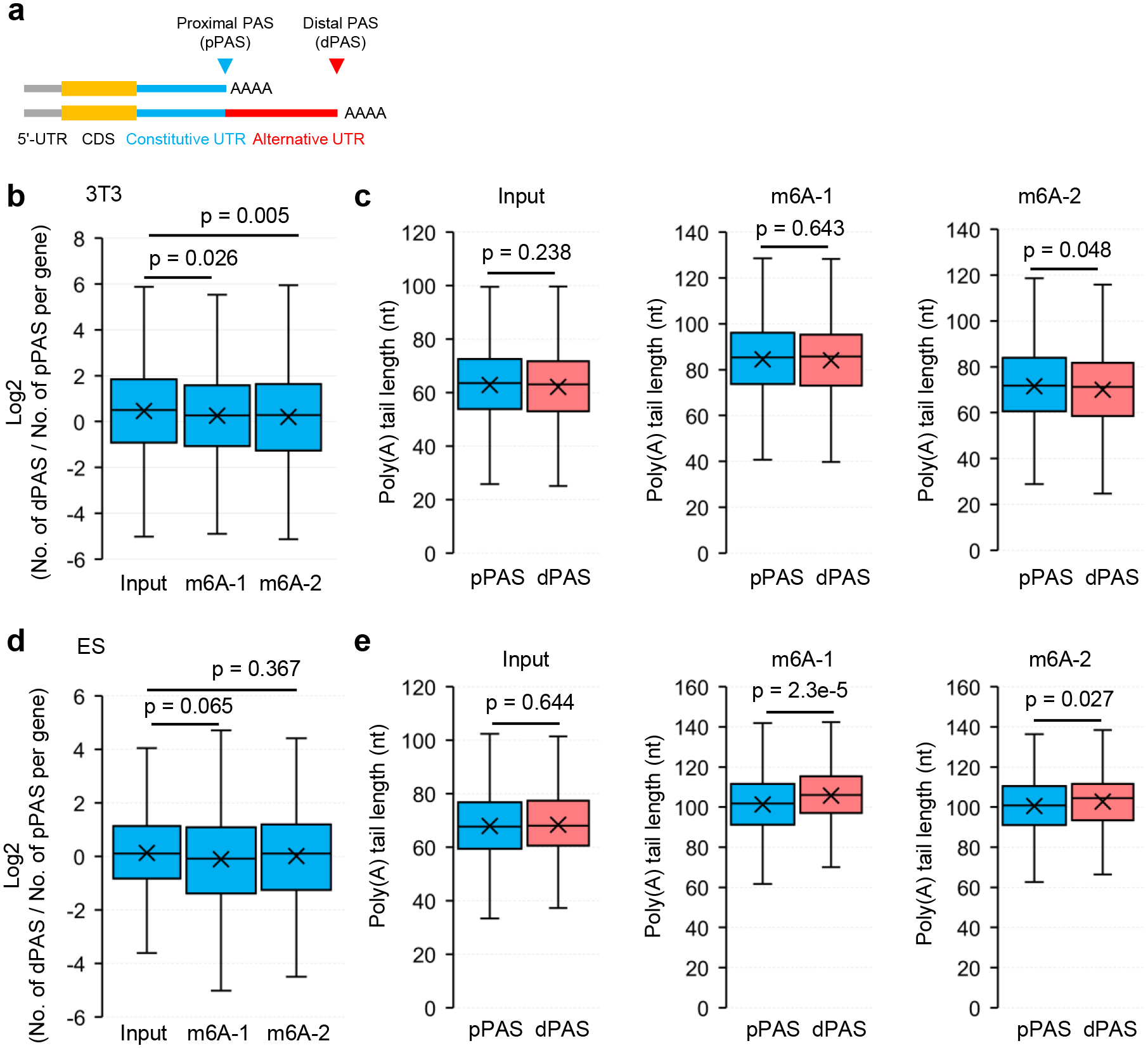
Relationship between m^6^A and alternative polyadenylation sites. **a,** Model showing genes with the proximal PAS (pPAS) and the distal PAS (dPAS). **b, d,** Box plot for the ratio of read counts between dPAS and pPAS isoforms per gene in log2 scale in m^6^A RIP and input samples of 3T3 (**b**) or ES (**d**). Genes (n = 1,021; 455 for 3T3 and ES, respectively) with at least 10 poly(A) tail containing reads (tail length ≥ 1 nt) for both dPAS and pPAS are included in the analysis. **c, e,** Box plot of poly(A) tail length per isoform using pPAS and dPAS in m^6^A RIP and input samples of 3T3 (**c**) or ES (**e**). Genes (n = 1,021; 455 for 3T3 and ES, respectively) with at least 10 poly(A) tail containing reads (tail length ≥ 1 nt) for both dPAS and pPAS are included in the analysis. *P* value is calculated using Student’s *t*-test. For all box plots, “×” indicates the mean value, horizontal bars show the median value, and the tops and bottoms of boxes represent the value of the 25^th^ and 75^th^ percentile, respectively.

These results suggest that cell type specific mechanisms exist in m^6^A preference or effect on alternative PAS choice and poly(A) tail length between transcript isoforms using different PAS.

### m^6^A-modified RNAs have longer poly(A) tails

Next, we investigated whether a relationship existed between m^6^A and poly(A) tail length. We calculated the proportion of transcripts containing poly(A) tails for each gene and found that m^6^A-modified transcripts, including mRNA and lncRNA, were more likely to have poly(A) tails in both 3T3 and ES (Fig. 4). A distribution map of poly(A) tail length for transcripts containing poly(A) tails was constructed and a statistically significant trend of m^6^A-modified transcripts, including mRNA and lncRNA, having longer poly(A) tails in both 3T3 and ES was observed (Fig. 4). Additionally, the median poly(A) tail length of each gene was compared in scatter plots and significantly longer poly(A) tails in m^6^A-modified transcripts in 3T3 and ES were found (Extended Data Fig. 3). These results indicate that m^6^A-modified transcripts possess longer poly(A) tails.

**Fig. 4.**
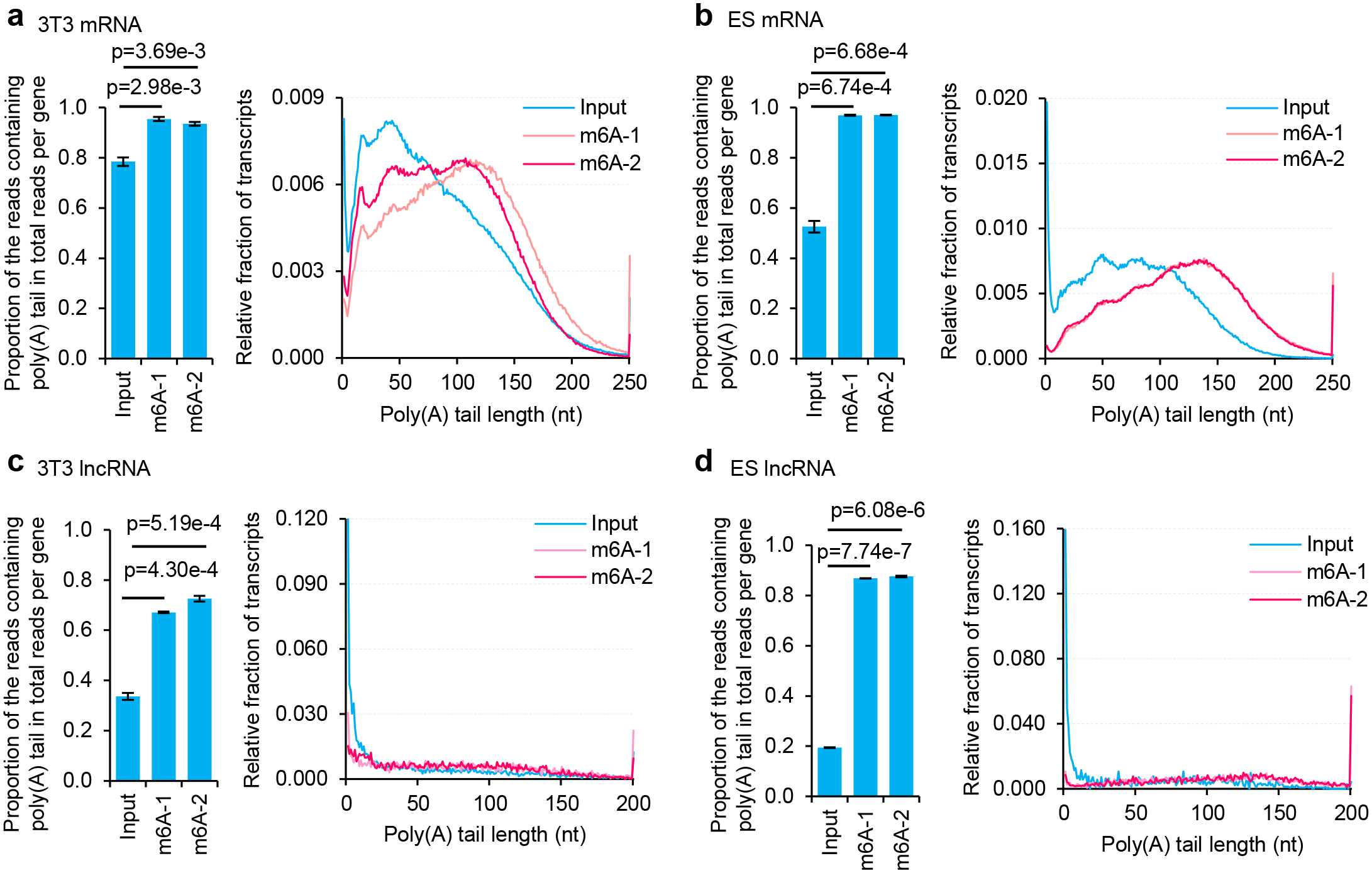
m^6^A-modified transcripts possess longer poly(A) tails. Left, Proportion of mRNAs (**a**, **b**) or lncRNAs (**c**, **d**) containing poly(A) tails in m^6^A RIP and input samples of 3T3 (**a**, **c**) or ES (**b**, **d**). Transcripts with poly(A) tails of at least 1 nt are included in the analysis. *p* value is calculated using Student’s *t*-test. Error bars indicate standard error of the mean (SEM) from two replicates. Right, Histogram of poly(A) tail length of mRNAs (**a**, **b**) or lncRNAs (**c**, **d**) in m^6^A RIP and input samples of 3T3 (**a**, **c**) or ES (**b**, **d**). Histograms (bin size=1 nt) are normalized to cover the same area. Transcripts with poly(A) tail of at least 1 nt are included in the analysis. Transcripts with poly(A) tail lengths greater than 250 nt (200 nt for **c**, **d**) are included in the 250 nt (200 nt for **c**, **d**) bin.

### The proportion of poly(A) tails containing non-A residues is lower for m^6^A-modified RNAs

Recently, non-A residues in poly(A) tails (non-A residues for short hereafter) were discovered and found to play roles in spermatogenesis^50^, oogenesis^51^, and maternal-to-zygotic transition^34–38^. This inspired investigation of whether m^6^A affects the incorporation of non-A residues in poly(A) tails. We analyzed the proportion of poly(A) tails containing non-A residues in m^6^A RIP and input samples of 3T3 and ES. We discovered that the proportion of mRNAs with non-A residues was less in m^6^A RIP than input samples in ES but not in 3T3 (Fig. 5a, b, left). The proportion of lncRNAs with non-A residues was less in m^6^A RIP than in input samples, both in ES and in 3T3 (Fig. 5c, d, left). m^6^A-modified transcripts, for both mRNAs (Fig. 5a, b, right) and lncRNAs (Fig. 5c, d, right), incorporated fewer non-A residues in the 5′-end, internal region, and 3′-end of poly(A) tails in both 3T3 and ES. At the gene level, m^6^A-modified transcripts incorporated fewer U and C into tails in 3T3 (Extended Data Fig. 4a), while m^6^A-modified transcripts incorporated much fewer U, C, and G into tails in ES (Extended Data Fig. 4b). These results indicate that the proportion of poly(A) tails containing non-A residues is lower for m^6^A-modified RNAs. Notably, we discovered that non-A residues are more enriched in shorter poly(A) tails in ES^39^. Considering that m^6^A-modified RNAs possess longer poly(A) tails (Fig. 4), we speculate this is one explanation for the proportion of transcripts containing non-A residues in their tails being lower for m^6^A-modified RNAs in ES cells.

**Fig. 5.**
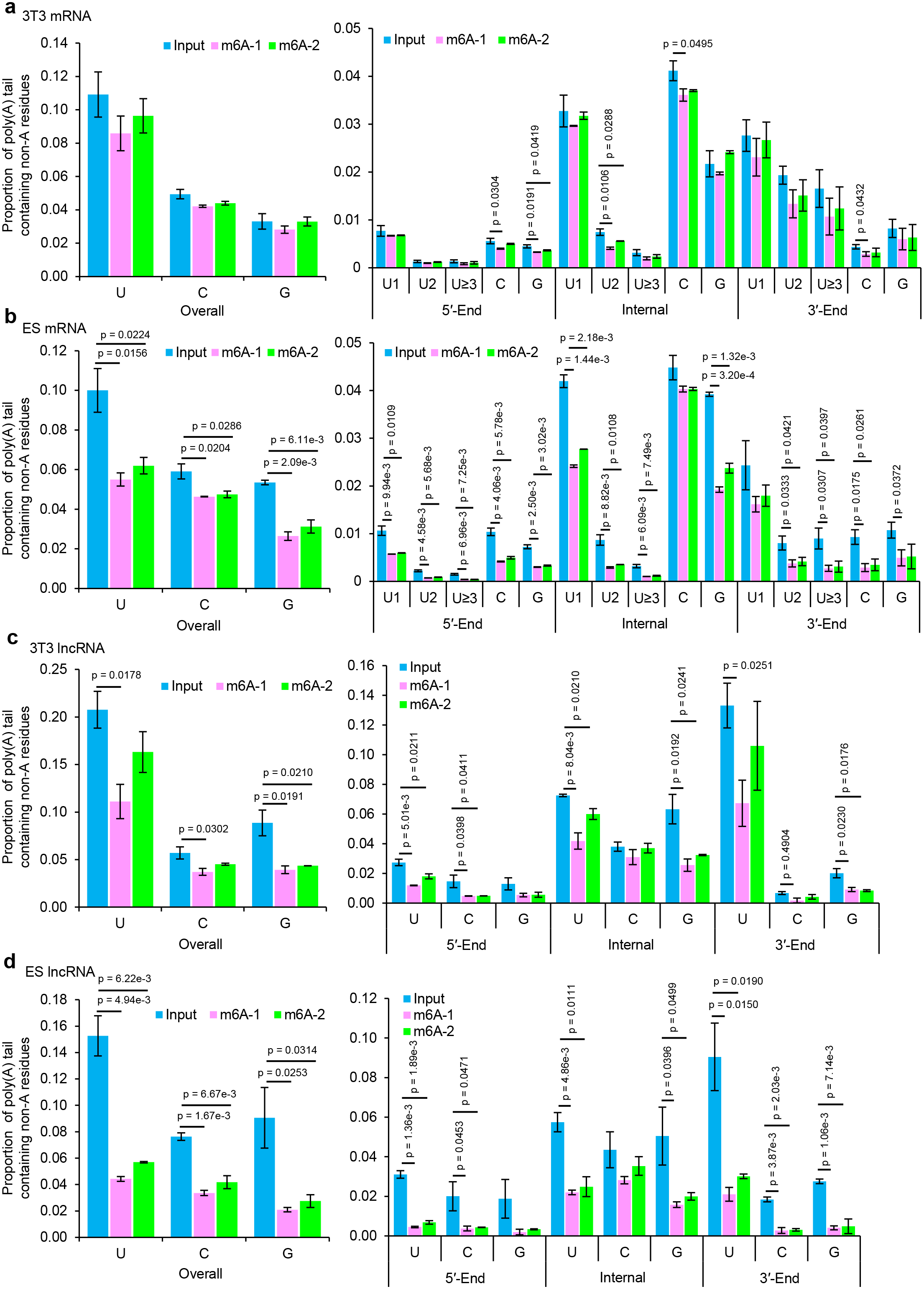
The proportion of poly(A) tails containing non-A residues is lower for m^6^A-modified RNAs. Proportion of mRNAs (**a**, **b**) or lncRNAs (**c**, **d**) containing non-A residues in the poly(A) tails in overall (left), or in the 5′-end, internal, and 3′-end of poly(A) tails (right) in m^6^A RIP and input samples of 3T3 (**a**, **c**) and ES (**b**, **d**). The U residues are further divided according to the length of the longest consecutive U sequence (U1, U2, and U≥3). Transcripts with poly(A) tails of at least 1 nt are included in the analysis. *P* value is calculated using Student’s *t*-test and is shown for groups that are statistically significant. Error bars indicate SEM from two replicates.

### Relationship between m^6^A level and gene expression level, poly(A) tail length, RNA half-life, or ribosome-binding efficiency

As the most abundant RNA chemical modification, there are contradictory reports about the role of m^6^A in regulating mRNA destabilization and translational enhancement^17, 18, 20, 52–54^. Previous analysis mainly focus on the association between number of m^6^A modified sites for a given gene and its mRNA half-life or translational efficiency^17, 18, 20, 52–54^. We have the transcript counts in both m^6^A RIP samples and input samples, therefore, we examined the relationship between m^6^A level (the ratio of the CPM in m^6^A RIP samples to the CPM in input samples, for each individual gene) and several indicators related to protein production efficiency, such as RNA expression level, poly(A) tail length, RNA half-life, and ribosome-binding efficiency.

No correlation was observed between m^6^A level and RNA expression level in both 3T3 and ES (Fig. 6a, e). Interestingly, we found moderate positive correlations between m^6^A level and poly(A) tail length in both 3T3 and ES (Fig. 6b, f), further confirming our observation of longer poly(A) tails in m^6^A modified transcripts. However, we found minimal, if any, correlation between m6A level and ribosome-binding efficiency^55, 56^ (Fig. 6c, g), or RNA half-life^46, 57^ (Fig. 6d, h), in both 3T3 and ES. These results indicate that m^6^A level is minimally correlated with gene expression level, RNA half-life, or ribosome-binding efficiency, but correlated with poly(A) tail length.

**Fig. 6.**
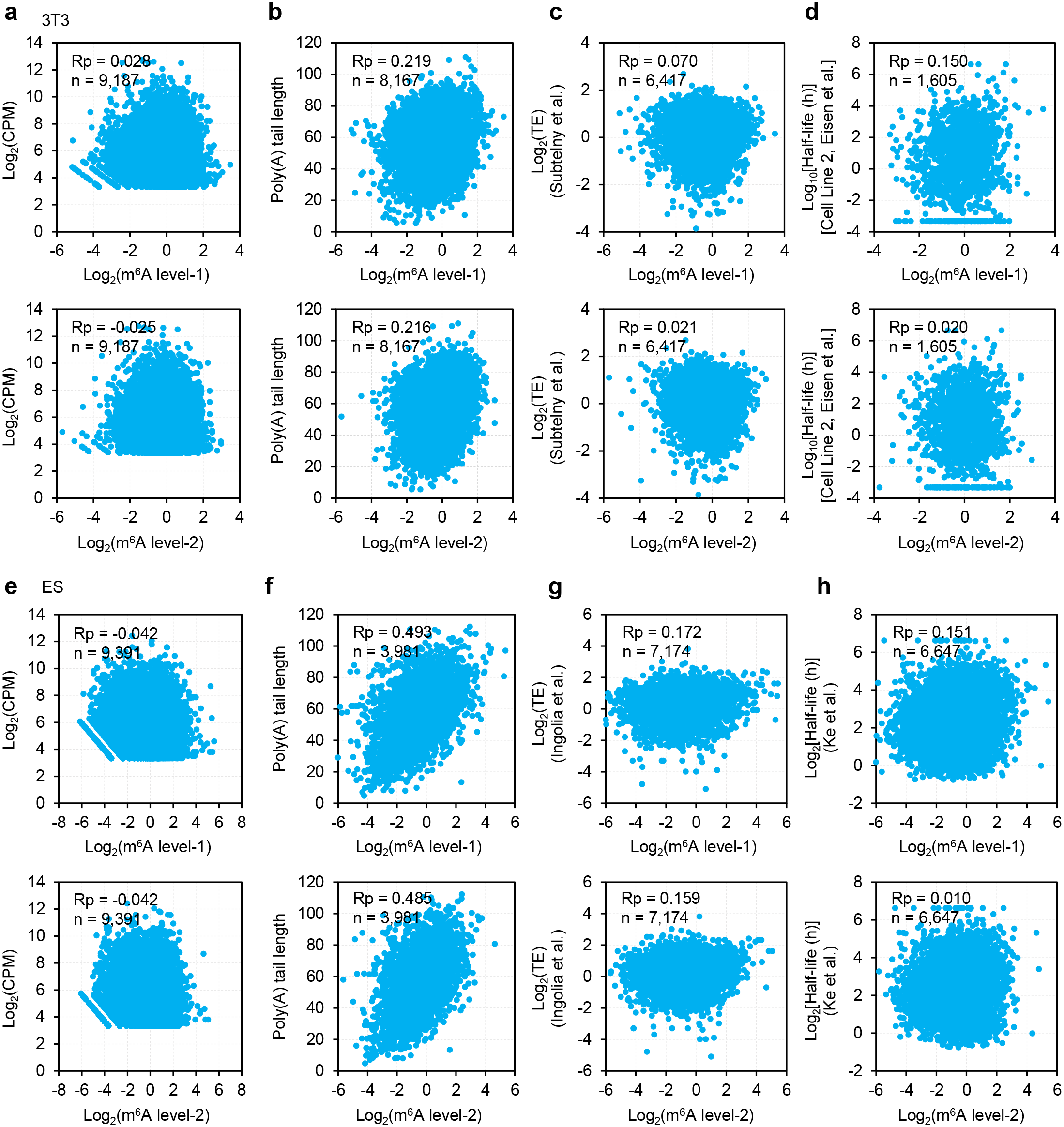
Relationship between m^6^A level and gene expression level, poly(A) tail length, RNA half-life, or ribosome-binding efficiency. **a, e,** Scatter plot of m^6^A level (ratio of CPM of m^6^A RIP samples to that of input, per gene, m^6^A level-1, Active Motif, #91261; m^6^A level-2, Abcam, #ab151230) and RNA expression level in log2 scale in 3T3 (**a**) and ES (**e**). **b, f,** Scatter plot of m^6^A level and poly(A) tail length in log2 scale in 3T3 (**b**) and ES (**f**). The poly(A) tail length for each gene is the geometric mean length of all the transcripts with poly(A) tails of at least 1 nt for the given gene. Genes with at least 20 reads in each sample are included in the analysis. **c, g,** Scatter plot of m^6^A level and translation efficiency (TE, ratios of CPM of ribosome-seq samples to CPM of input, per gene) in log2 scale in 3T3^55^ (**c**) and ES^56^ (**g**). **d, h,** Scatter plot of m^6^A level and RNA half-life in log2 scale in 3T3^57^ (**d**) and ES^46^ (**h**). For all scatter plots, each dot represents one gene. Pearson’s correlation coefficient (Rp) and number of genes included in the analysis are shown at the top left of each graph.

## Discussion

Current m^6^A annotation methods such as MeRIP-Seq^58^ are mainly based on second-generation short-reads sequencing technologies, which cannot allow a full view of the full-length transcripts and poly(A) tails. In this study we developed a new method, MePAIso-seq2, with the ability to sequence poly(A) tail inclusive full-length m^6^A modified transcripts, to study the relationship between m^6^A, alternative splicing and poly(A) tailing. Using MePAIso-seq2, we clearly demonstrate that m^6^A promotes and inhibits similar numbers of AS events in 3T3 and mouse ES cells, but has no obvious effect on alternative PAS choice, which differs from previous reports^46, 47, 49^. Interestingly, we discovered that m^6^A-modified transcripts possess longer poly(A) tails. Additionally, we found the ratio of poly(A) tails containing non-A residues was lower in m^6^A modified transcripts, especially in ES (Fig. 7).

**Fig. 7.**
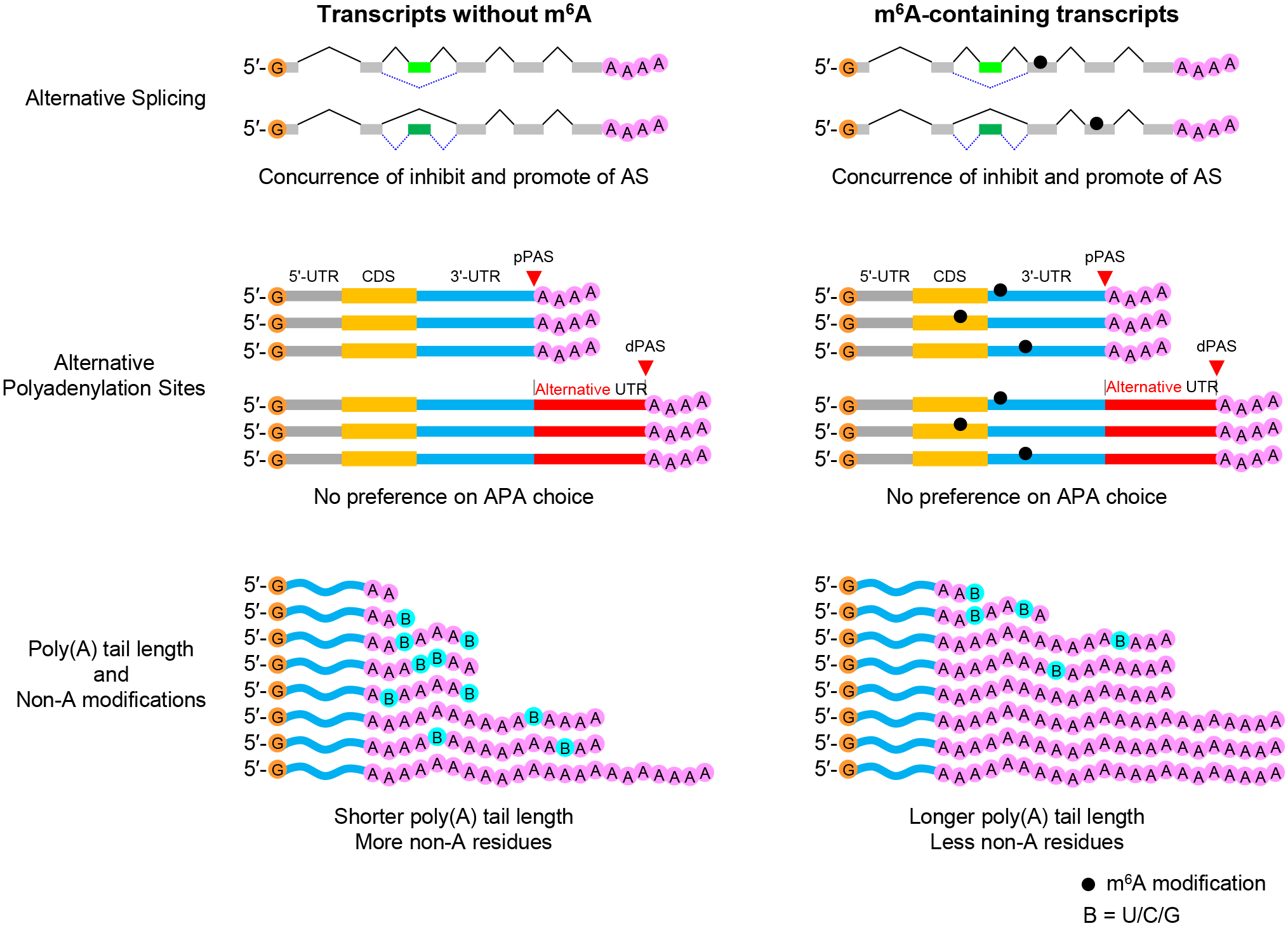
Model for the relationship between m^6^A and alternative splicing, PAS choice, poly(A) tail length, and non-A residues. The model shows that m^6^A regulates RNA splicing but has no effect on PAS choice, and m^6^A-modified transcripts possess longer poly(A) tails and with lower proportion of poly(A) tails containing non-A residues.

One possible reason for the lower level of poly(A) tails containing non-A residues is that transcripts in m^6^A RIP samples possess longer poly(A) tail. We previously reported that non-A residuals are more enriched in shorter poly(A) tails in ES^39^. Therefore, the longer poly(A) tail and lower level of poly(A) tails containing non-A residues in m^6^A modified transcripts are likely results of a common regulatory mechanism controlling the polyadenylation of the m^6^A modified transcripts. This mechanism is of great interest and warrants future investigation. The biological functions of m^6^A are mediated through its “readers” such as the YT521-B homology (YTH) domain family of proteins (YTHDF1, YTHDF2, YTHDF3 and YTHDC1)^5, 59, 60^, and Insulin-like growth factor-2 mRNA-binding proteins (IGF2BP1, IGF2BP2, and IGF2BP3)^42^. Thus, it is worthwhile to study whether these m^6^A “reader” proteins interact with poly(A) tail processing proteins such as poly(A)-binding proteins (PABPs) as well as multiple PABP-interacting factors.

Minimal correlation was found between m^6^A level and RNA half-lives, or ribosome-binding efficiency in general. Previous publications have contradictory reports on the role of m^6^A on mRNA half-life and translational efficiency^17, 18, 20, 52–54^. One central debate is whether the binding of different m^6^A reader proteins to the m^6^A modified mRNA leads to the same or different outcome. Some report suggest that different m^6^A reader proteins binds different set of m^6^A modified mRNA and leads to reader specific outcome^17, 18, 20, 53, 54^. However, a recent report propose that the different reader proteins are of similar m^6^A binding properties and combination actions of these readers lead to degradation of the modified mRNA^52^. Our data focus on the m^6^A modifications themselves which does not involve the reader proteins. Therefore, it is reasonable that our observations might be different from those reports based on data from the manipulation of the readers. In addition, different from the previous analysis focusing on the presence of m^6^A sites in the gene or not, our data provide a different aspect in considering the actual level of m^6^A modification for each gene rather than merely the numbers of m^6^A modification site. Given that the actual modification level information in our study, we believe that our data will help in the study of the biological outcome of m^6^A modifications in the future by considering the actual level of m^6^A modification beyond just presence of not.

RNA modifications, including structural modifications (cap and poly(A) tail) and chemical modifications (over 170 types are identified, like m^6^A and N4-acetylcytidine (ac4C)), expand the RNA code and give rise to a new field called “RNA epigenetics”^3, 61^. It is becoming increasingly interesting to study cross-talks between chemical modifications and the structural poly(A) tails. Whether these RNA modifications cooperate or compete to alter the metabolism and function of RNAs post-transcriptionally warrants further investigations. Our newly developed MePAIso-seq2 method makes it possible to dissect the interactions between these modifications, which will shed new lights on their roles in diverse physiological and pathological processes.

## Material and Methods

### Culture cells

NIH 3T3 cells (3T3) were grown in DMEM (Invitrogen, 12100-046) supplemented with 10% FBS (Gibco, 16140071) and 1 mg/mL puromycin, and incubated at 37 ℃ in 5% CO_2_. Mouse embryonic stem cells (ES, E14) were cultured without feeder cells in DMEM (Invitrogen, 12100-046) containing 15% FBS (Gibco, 16140071), leukemia inhibiting factor (LIF, Sigma, SRP6287), penicillin/streptomycin (Hyclone, sv30010), L-glutamine (Gibco, 25030081), β-mercaptoethanol (Sigma, M3148), and non-essential amino acids (Gibco, 11140-050), and incubated at 37 ℃ in 5% CO_2_. Both 3T3 and ES cells were free of mycoplasma contamination.

### Total RNA isolation and m^6^A RNA immunoprecipitation (RIP)

Total RNA was extracted from cells using TRIzol reagent (Invitrogen, 15596018) according to the manufacturer’s instructions and stored at −80 °C or used immediately. Total RNA samples were subjected to immunoprecipitation with m^6^A specific antibody (m^6^A-1, Active Motif, #91261; m^6^A-2, Abcam, #ab151230) as previously described^5^. Briefly, the immunoprecipitation (IP) reaction mixtures containing total RNA, m^6^A specific antibody, Ribonucleoside vanadyl complexes (Sigma, R3380) and RNasin Plus RNase inhibitor (Promega, N2611) in IP buffer^5^ were prepared and incubated with head-over-tail rotation for 2 hours at 4 °C. After adding protein A beads, the reaction mixtures were incubated for another 2 hours on a rotating wheel at 4 °C. After washing, m^6^A-modified RNAs were eluted from beads and used for library preparation and sequencing.

### PAIso-seq2 library construction and data pre-processing

PAIso-seq2 libraries were constructed following the PAIso-seq2 protocol described in another study^39^. The PAIso-seq2 data generated in this study was processed following the methods described^35, 37^. Clean CCS reads were aligned to the *Mus musculus* UCSCmm10 reference genome using *minimap2* (v.217-r941). The poly(A) tail sequences were extracted using python script PolyA_trim.py (http://). The 3′-soft clip sequence of CCS reads in the alignment file were used as candidate poly(A) tail sequences. The 3′-soft clip sequences with the frequency of U, C, and G greater or equal to 0.1 simultaneously were marked as “HIGH_TCG” tails. To better define the poly(A) tail, we defined a continuous score based on the transitions between the two adjacent nucleotide residues throughout the 3′-soft clip sequences. To calculate the continuous score a transition from one residue to the same residue scored 0, while a transition from one residue to a different residue scored 1. The 3′-soft clips which were not marked as “HIGH_TCG” and with continuous scores less than or equal to 12 were considered as poly(A) tails. The U, C, and G residues in the poly(A) tail are considered non-A residues.

### Alternative splicing analysis

Seven types of alternative splicing (AS) events, including alternative 3′ splice site (A3), alternative 5′ splice site (A5), alternative first exon (AF), alternative last exon (AL), mutually exclusive exon (MX), retained intron (RI), and skipped exon (SE) were identified using *SUPPA2* (v2.2.1)^62^. Genes associated with AS events were defined as AS genes. *SUPPA2* was also used to calculate the percent spliced in index (PSI) of each AS event using default parameters. ΔPSI was calculated as the PSI of input sample minus PSI of m^6^A RIP sample for each AS event. The ΔPSI and its associated *p*-*value* between input and m^6^A RIP samples were calculated using the *diffSplice* method in the *SUPPA2* package. AS events with ΔPSI < 0 and *p < 0.05* were treated as m^6^A-promotion events; AS events with ΔPSI > 0 and *p < 0.05* were treated as m^6^A-inhibition events.

### Polyadenylation site calling for annotated mRNA

For each type of cells, the input PAIso-seq2 data were used for polyadenylation site calling. Clean CCS reads were aligned to the reference genomes using *minimap2*^63^. A greedy strategy was used for computing the depth of each candidate poly(A) site (namely the 3′-end of the alignment) as the number of reads aligned within a window of 5 nucleotides of the site. The site with maximum read depth across all candidates was added to the list of poly(A) sites if the site had a depth of at least 10 and did not occur within 20 nucleotides of a previously added site. The site was then removed from the list of candidates and the algorithm proceeded until no candidate sites remained. Protein-coding genes in the nuclear genome were included in this analysis.

### Classification of APA isoforms for each gene

For each cell type, the polyadenylation sites called in the input data were used as reference point for classification of APA isoforms. Genes with two or more reference polyadenylation sites were included in the analysis. Transcripts with polyadenylation sites located within 5 nt of the reference polyadenylation sites were classified as the transcripts associated with the given polyadenylation site. Protein-coding genes in the nuclear genome were included in this analysis.

### Data availability

The ccs data in bam format from MePAIso-seq2 and PAIso-seq2 experiments will be available at Genome Sequence Archive hosted by National Genomic Data Center. This study includes analysis of the following published data: the m^6^A-RIP-RNA-seq data of Huang et al. (Gene Expression Omnibus database (GEO) accession no. GSE90639) and Wen et al. (GEO accession no. GSE94148) used in Fig. 1c; ribosome profiling data of Subtelny et al. (GEO accession no. GSE52809) used in Fig. 6c and Ingolia et al. (GEO accession no. GSE30839) used in Fig. 6g; and RNA half-life data of Eisen et al. (GEO accession no. GSE134660) used in Fig. 6d and Ke et al. (GEO accession no. GSE86336) used in Fig. 6h. Custom scripts used for data analysis will be available upon request.

## Acknowledgements

We thank Ying Liu for her technical assistance in cell culturing. We thank Hu Nie for his technical assistance in bioinformatic analysis. This work was supported by the National Key Research and Development Program of China (2020YFA0804000), the Strategic Priority Research Program of the Chinese Academy of Sciences (XDA24020203), National Natural Science Foundation of China (31970588, 32170606), Natural Science Foundation of Heilongjiang province (YQ2020C003), the China Postdoctoral Science Foundation (2020M670516, 2020T130687), and the State Key Laboratory of Molecular Developmental Biology.

## Author Contributions

Yusheng Liu, Falong Lu and Jiaqiang Wang conceived the project and designed the study. Yusheng Liu constructed the library of the MePAIso-seq2. Yusheng Liu, Yiwei Zhang, Falong Lu and Jiaqiang Wang analyzed the sequencing data. Yusheng Liu and Jiaqiang Wang organized all figures. Yusheng Liu, Falong Lu and Jiaqiang Wang supervised the project. Yusheng Liu, Falong Lu and Jiaqiang Wang wrote the manuscript with the input from the other authors.

## Competing Interests statement

The authors declare no competing interests.

---

**Extended Data Fig. 1.**
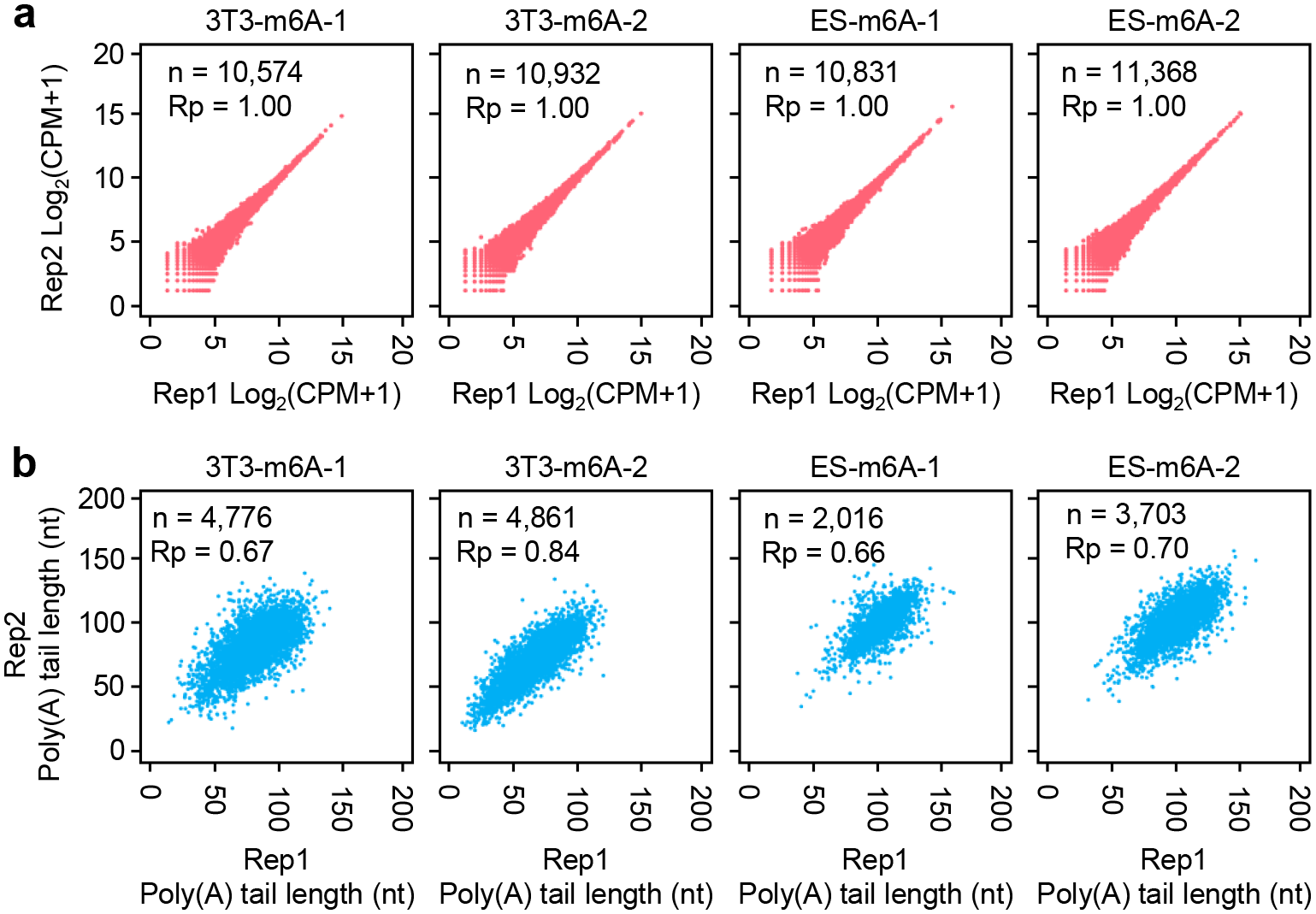
The reproducibility of MePAIso-seq2 data. **a,** Scatter plot showing the Pearson correlation of gene expression between two replicates for MePAIso-seq2 in 3T3 and ES. Each dot represents one gene. Pearson’s correlation coefficient (Rp) and number of genes included in the analysis are shown at the top left of each graph. **b,** Scatter plot showing the Pearson correlation of poly(A) tail length between two replicates for MePAIso-seq2 in 3T3 and ES. Each dot represents one gene. The poly(A) tail length for each gene is the geometric mean length of all the transcripts with poly(A) tails of at least 1 nt for the given gene. Genes with at least 20 reads in each sample are included in the analysis. Pearson’s correlation coefficient (Rp) and number of genes included in the analysis are shown at the top left of each graph.

**Extended Data Fig. 2.**
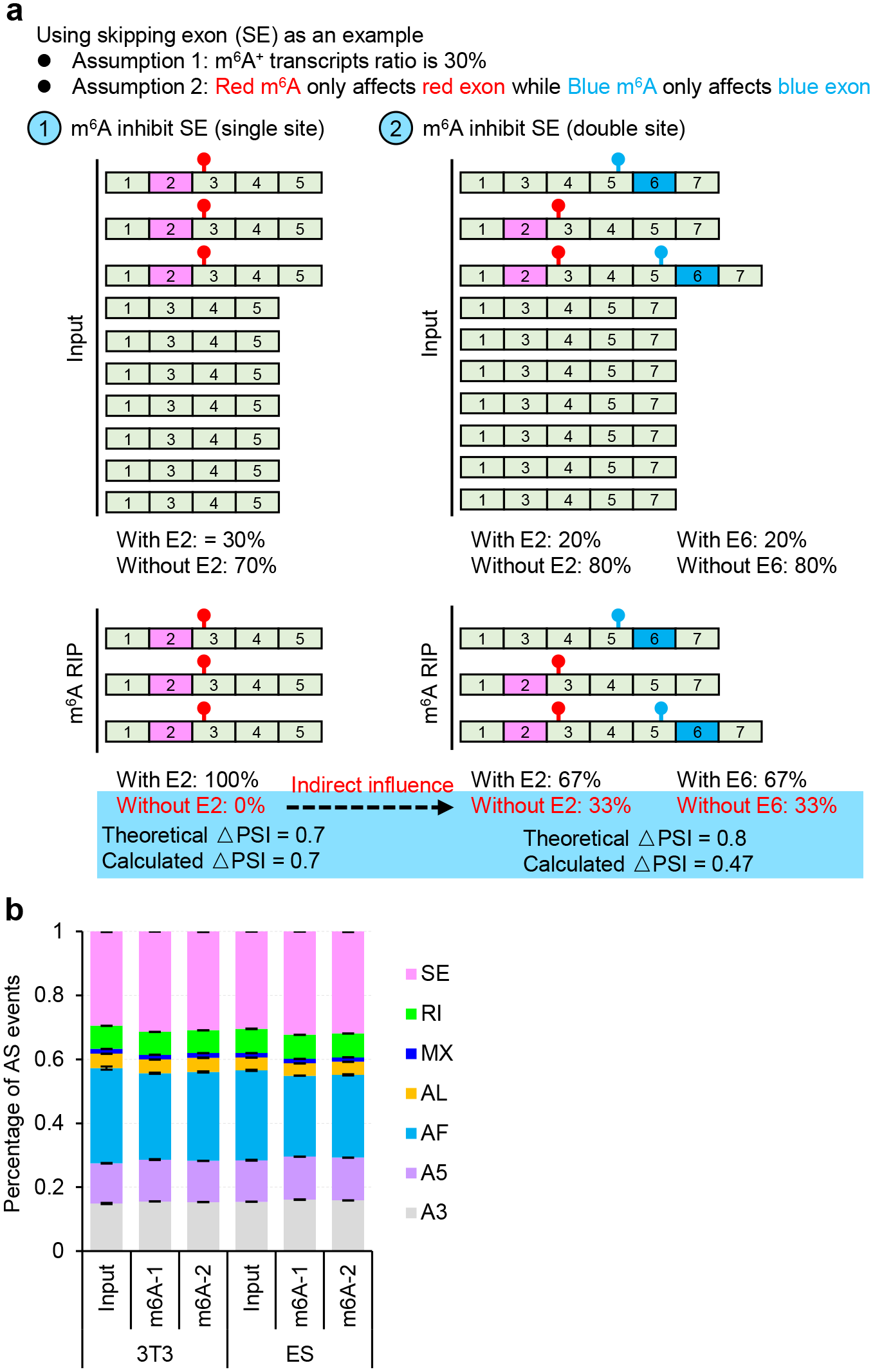
m^6^A regulates mRNA splicing without bias in AS types. Model showing the calculated ΔPSI must be less than the theoretical value. If there is only one m^6^A and one AS event in each transcript, the calculated ΔPSI must be equal to the theoretical value, however, if the number of m^6^A sites and the AS events in one transcript are more than one, the calculated ΔPSI must be less than the theoretical value, because each m^6^A may not affect all the AS events. Proportions of the AS events of the seven types identified from m^6^A RIP and input samples. AS, alternative splicing; A5, alternative 5′ splice site; A3, alternative 3′ splice site; AF, alternative first exon; AL, alternative last exon; MX, mutually exclusive exon; RI, retained intron; SE, skipped exon. Error bars indicate standard error of the mean (SEM) from two replicates.

**Extended Data Fig. 3.**
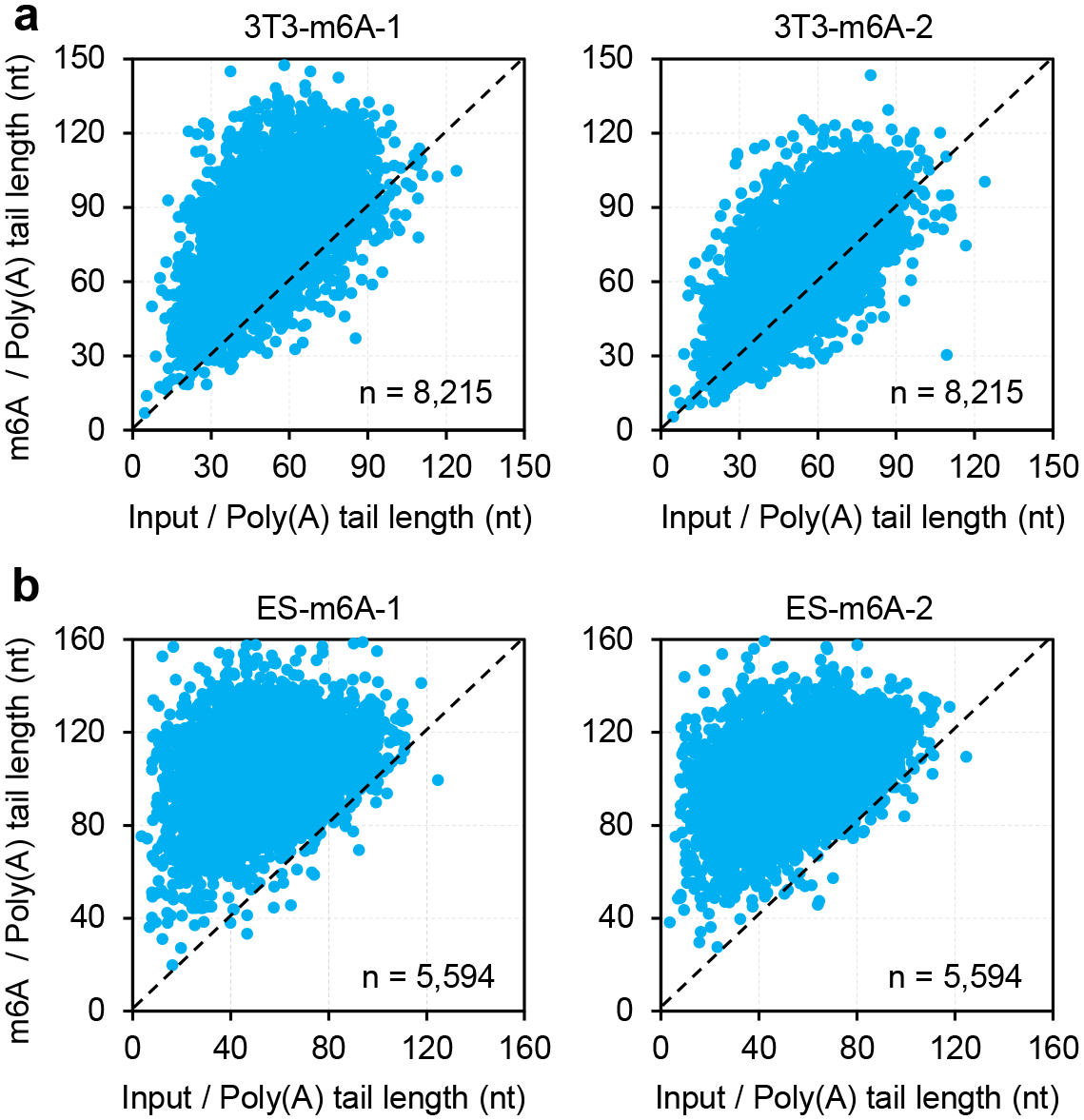
m^6^A-modified transcripts own longer poly(A) tails. Scatter plot of poly(A) tail length of m^6^A RIP and input samples of 3T3 (**a**) and ES (**b**). Each dot represents one annotated coding gene. The poly(A) tail length for each gene is the geometric mean length of all transcripts with poly(A) tails of at least 1 nt for the given gene. Genes with at least 10 reads in each sample are included in the analysis. Number of genes included in the analysis is shown at the bottom right of each graph.

**Extended Data Fig. 4.**
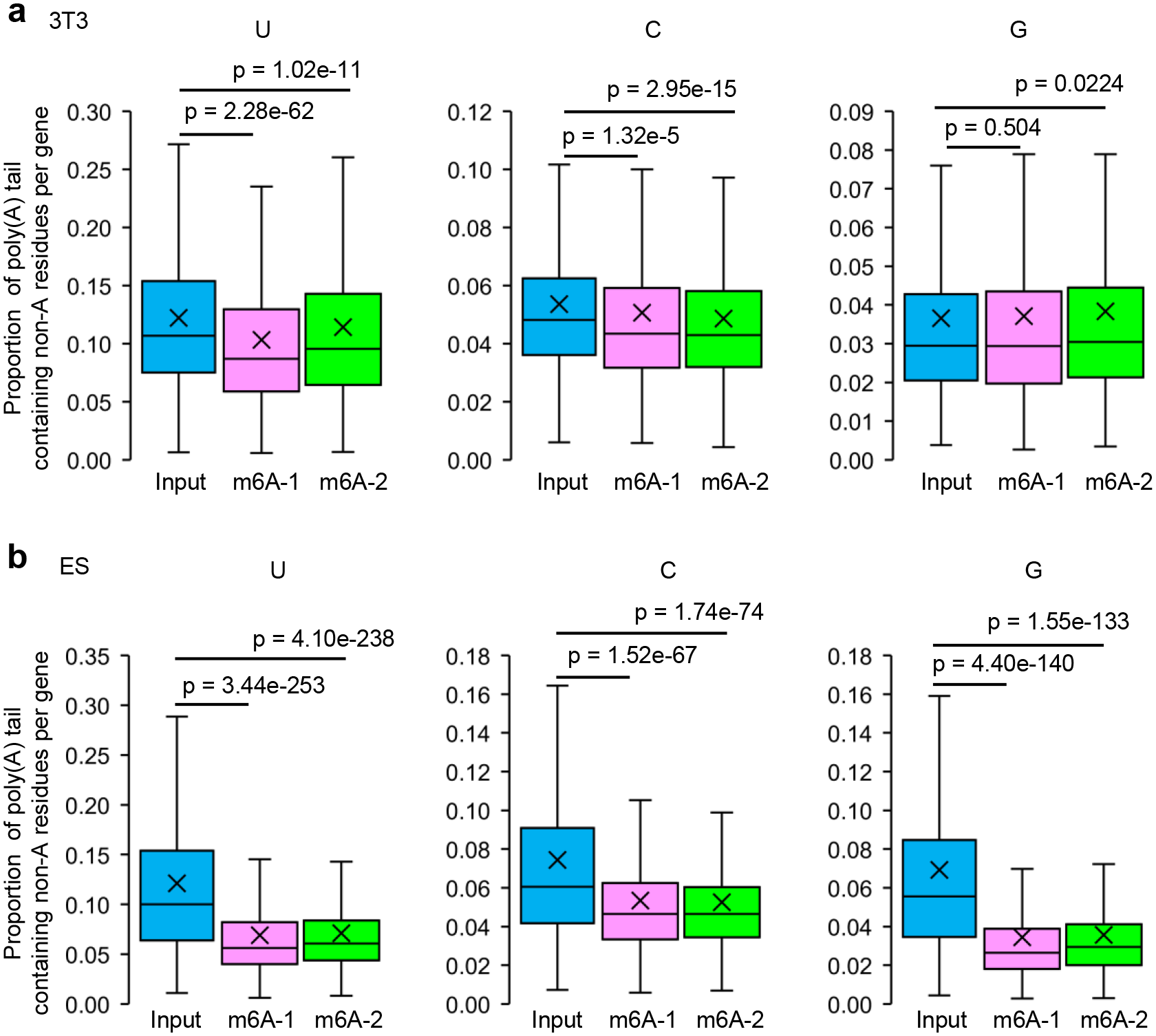
m^6^A-modified transcripts incorporate less non-A residues. Box plot for proportion of transcripts containing non-A residues in tails for genes in m^6^A RIP and input samples of 3T3 (**a**) and ES (**b**). Annotated coding genes (gene numbers: 3T3-U, 7264; 3T3-C, 5791; 3T3-G, 4585; ES-U, 4322; ES-C, 3562; ES-G, 2821) with at least 10 reads and poly(A) tails of at least 1 nt are included in the analysis. For all box plots, “×” indicates the mean value, black horizontal bars show the median value, and the tops and bottoms of the boxes represent the value of the 25^th^ and 75^th^ percentile, respectively. *P*-value is calculated using Student’s *t*-test.

## Notes

### Competing Interest Statement

The authors have declared no competing interest.

